# Biophysical Design Space for Cellular Self-assembly and Dynamics

**DOI:** 10.1101/2025.11.13.688226

**Authors:** Sanbed Das, M. Sreepadmanabh, Dikshant Parashar, Tapomoy Bhattacharjee, Sayantan Dutta

## Abstract

In natural biological systems, cells organize into tissues through interactions of several processes, including cellular signaling, collective migration, contractile activity of cytoskeletal elements and interactions with their surroundings. In recent decades, advancements in microscopy, genetic engineering, biochemistry, and computational modeling have enabled a more quantitative understanding of these processes. In this article, we present an integrated computational framework that couples various physical mechanisms such as cell-cell adhesion, strength and persistence of cellular motility, and the background stiffness. We use this framework to study how they collectively interact to determine the self-organization of the cells from a pseudo-random structure as well as the migration behavior of the cells. Notably, our simulations predict that motility has a two-way effect on cellular self-assembly: it promotes aggregation at moderate levels but disrupts clusters when excessively strong, yielding an optimal motility for formation of multicellular clusters. On the other hand, adhesion shows a two-stage effect: At lower value it self-assembles the structure, at higher value it compacts it. Furthermore, we experimentally demonstrate the *motility-assisted* self-aggregation of cells using cancer cells in a granular mechanical milieu. Finally we show that cell-cell adhesion and background medium tune the strength and persistence of cellular migration. Altogether, this work presents a computational framework that allows us to design phase behavior of collective of cells tuning their interaction, motility, and the background mechanics.

## 1 Introduction

Cells, fundamental building blocks of life, self-assemble to spatially organized and functional structures such as organs and tissues. Self-organization of cells is a central *design* principle in numerous physiological processes, including tissue pattern formation leading to embryonic morphogenesis [1, 2], wound healing [3–5], immune response [6], and regeneration. When these *designs* fail, consequences can be severe, leading to pathological conditions such as tumor invasion [7] or fatal developmental abnormalities [8, 9]. The same organizing principles are often leveraged in generating *in vitro* assemblies of cells such as organoid [10]. A set of biophysical mechanisms interact with each other and set the *design* rules of this self-organization. Cells bind to one another through E-cadherin–mediated adhesion [11] and form multicellular clusters [12]. As they proliferate, local density increases and cells generate specific packing to accomodate each other [13, 14]. Cells migrate—sometimes directionally following a cue (taxis) [15, 16], sometimes randomly (kinesis) [17] to explore the surrounding medium. Cell-cell communication occurs through chemical signaling as well as through forces transmitted by cytoskeletal elements such as microtubules [18] and actomyosin [19, 20]. At the same time, cells interact with the biochemical and mechanical properties of their surrounding medium and respond passively or adjust their behavior [21, 22]. Together, these mechanisms promote the spontaneous formation of organized multicellular assemblies with specific structure and function.

Active matter physics has devoted considerable effort over the last decade to modeling the self-organization and dynamics of self-propelled particles [23–25]. Unlike passive colloidal particles, whose aggregation is governed by equilibrium forces, self-propelled particles can reorganize even without attractive interactions. A key example is motility-induced phase separation, where self-propulsion alone drives the formation of dense clusters and spatially segregated domains [26, 27]. Typical ingredients used to model the self-organization of such particles include self-propulsion described as Ornstein–Uhlenbeck noise [28] or active Brownian motion [29], short range inter-particle forces [29–31], alignment of propulsion directions between neighboring units captured by the Vicsek model [32], and stiffness of the particles [33]. These interactions generate a rich set of emergent behaviors, including dynamic clustering [34, 35], flocking [36–39], and swarming [40] and transitions between jammed to unjammed states [41].

Physical realizations of self-propelled interacting active matter span multiple scales, from reactive droplets [42] and biomolecular condensates at molecular dimensions [43] to insect swarms and schools of fish at population scales [44, 45]. The self-organization of cells are also examples of such system in an intermediate lengthscale as cells are interacting, self-propelled active particles [46–48]. In this article, we present a computational framework that describes the self-assembly of cells using principles from soft and active matter physics. Specifically, we treat cells as soft self-propelling particles that adhere to one another and are submerged in a viscoelastic medium represented by gels, and using this framework we demonstrate how different biophysical mechanisms interplay with each other to determine the cellular self-assembly and dynamics. Specifically, we show that motility has a two-way effect in self-assembly by facilitating cell-cell assembly on one hand and destabilizing already assembled clusters on the other hand, leading to an optimal activity window for robust multicellular organization. In contrast, cell–cell adhesion exhibits a two-stage influence: weak adhesion enables formation of cluster, while stronger adhesion primarily drives compaction. We further provide experimental evidence of this *motility-assisted* aggregation using cancer cells suspended in a granular mechanical environment. Finally, we demonstrate that the interplay between cell–cell adhesion and the mechanical properties of the surrounding medium regulates both the strength and the persistence of cellular migration.

## 2 Model Description

We employ a particle-based simulation framework to investigate the dynamics of cellular self-organization. Our model consists of two kinds of particles: cells and background particles (denoted as gels) representing a viscoelastic medium. The choice of background material is motivated by hydrogel mediums used for cell-culture and 3D bioprinting in deformable medium [49, 50]. We perform the simulations in a square (2D)/cubic (3D) box with periodic boundary conditions [Fig. 1 (a),(b)]. The equation of motion for a particle *i* follows overdamped Langevin dynamics:

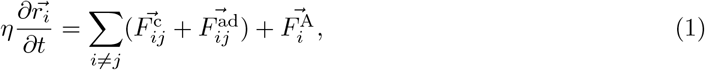

where, *η* is the coefficient of viscous drag,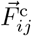, and 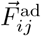 represent the contact and adhesive force from all other particles *j*, and 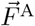 represents the active force resulting into the motility.

**Fig. 1.**
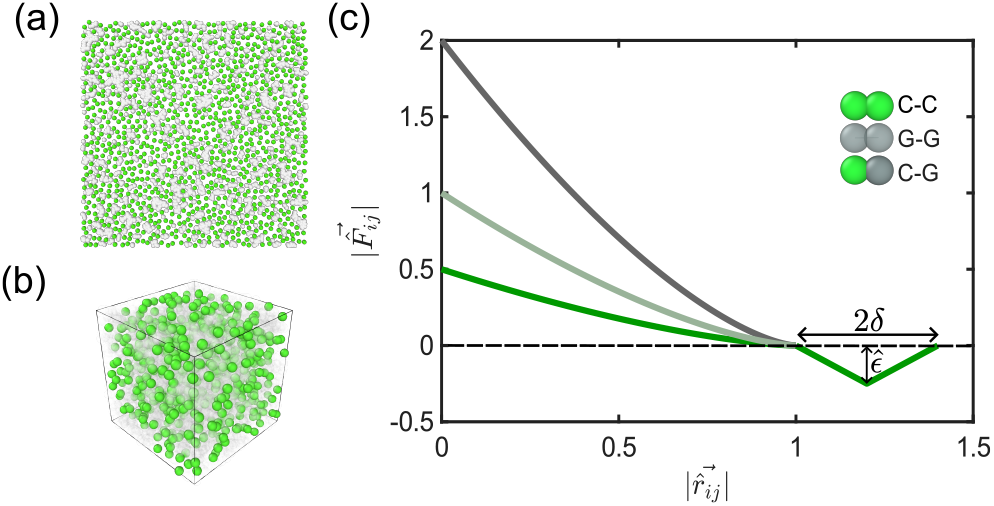
Simulation snapshots from (a) two and (b) three dimensional simulation boxes respectively. Green particles denote cells and gray particles denote background (gel) particles. Gel particles are made transparent in the 3D simulation box for better visibility of cells. (c) The signed magnitude of inter-particle interaction forces 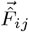 as a function of the radial center-to-center distance 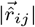. The positive values denote repulsion and the negative values represent attraction. The green, light-gray, and dark-gray curves represent cell-cell, cell-gel and gel-gel interactions.

The pairwise interaction forces consist of the contact force 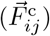 and the adhesion force 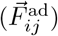. Hertz force [51] is often used to describe the contact interaction between soft particles specifically for microgels and cells [52, 53] and we adapt that to describe the contact interaction [Fig. 1(c)]:

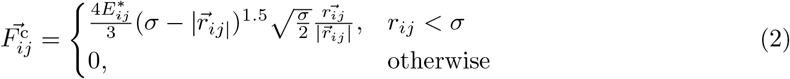

where, 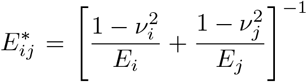 [54]. *E*_*i*_ and *E*_*j*_ are the Young’s moduli and *v*_*i*_ and *v*_*j*_ are the Poisson’s ratio of the particles *i* and *j*. All particles have the same diameter *σ*, and 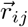 is the center-to-center separation vector. Equation 2 defines a soft repulsive force that acts only when particles overlap.

Next, we non-dimensionalize the length by the diameter of the particle σ and time by 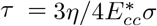, the viscoelastic relaxation timescale associated with the elastic modulus of the cells and viscosity of the background medium. The non-dimensional equation of motion is:

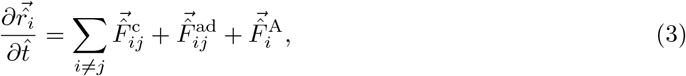

where, 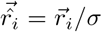 and 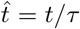. As a result, the non-dimensional forces are normalized by a factor of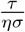. Specifically, the non-dimensional contact force reads as:

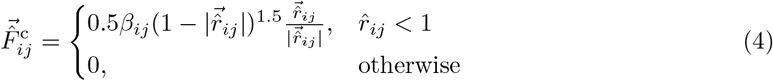

where, *β* _*ij*_ = 1 for cell-cell contact, 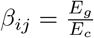 gel-gel contact and 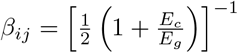 for cell-gel contact. We assume the Poisson’s ratio to be same for all the particles.

Cell-cell adhesions are often represented by short-range attraction force similar to a Lennard-Jones interaction with a finite-cutoff [2]. We specifically use a V-shaped force that acts only between pairs of cells[Fig. 1(c)]:

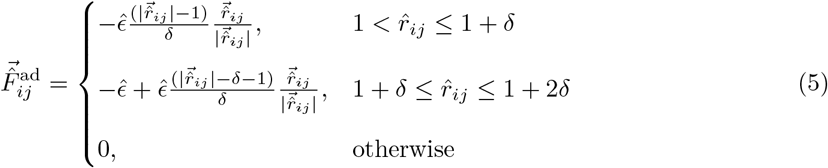

where, 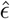 represents the strength of adhesion, and δ represents the half-width of the range of attractive force.

The active force in our model represents cellular motility and is inspired from the Ornstein-Uhlenbeck process [55, 56]. Active force on cell *i* evolves as,

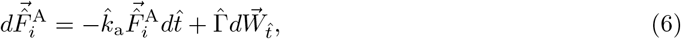

where, 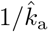 represents the timescale of persistence of the cellular motion, and 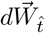 represents a Gaussian white noise, and 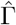 represents the strength of cellular motility. Following the properties of an Ornstein-Uhlenbeck process, the time correlation of the force evolves as,

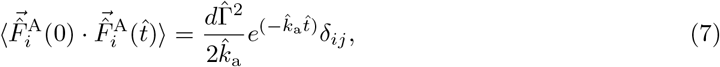

where, *d* is the dimension of the simulation box.

Altogether, we explore the effects of the following parameters in our simulation: 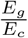 (relative stiffness of the background medium and the cells), 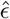 (the strength of adhesion), 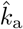 (inverse persistence timescale), 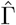 (the strength of motility), and *ϕ*_*c*_ (packing fraction of the cells). In this study we use a densely packed box, where the volume of undeformed cells and gels are equal to the volume of the box (i.e. *ϕ*_*c*_ + *ϕ*_*g*_ = 1, where *ϕ*_*g*_ is the volume fraction of the gels). Specifically, we create a non-overlapping configuration of cells using Random Sequential Addition (RSA)[57] and fill the rest of the space with gels, which may overlap with each other but don’t overlap with the cells.

## 3 Results and Discussions

In this section of the manuscript, we examine how the physical parameters introduced above influence collective cell self-assembly, specifically focusing on the formation of multicellular clusters, as well as the dynamics of individual cells.

### 3.1 Quantifying and mapping the degree of self-assembly

To quantify the degree of self-assembly, we identify clusters of connected cells. Two cells are defined as connected if their separation is less than a cutoff radius *r*_c_ = (1 + *δ*)*σ* (Section 5.1.2 of Methods), where *δ* is the non-dimesionalized half-width of the range of the attractive potential [Fig. 1(c)] and *σ* is the cell diameter. Based on this criterion, we construct an adjacency matrix and determine the number of independent clusters, *N*_conn_. We define the order parameter *S* = *N*_conn_*/N*_cell_, where, *N*_cell_ is the total number of cells [Fig. 2(a)]. *S* measures the degree of self-assembly. *S* = 1 corresponds to fully isolated cells, while *S* = 1*/N*_cell_ ≈ 0 corresponds to a single connected cluster. Thus, lower *S* indicates stronger self-assembly.

**Fig. 2.**
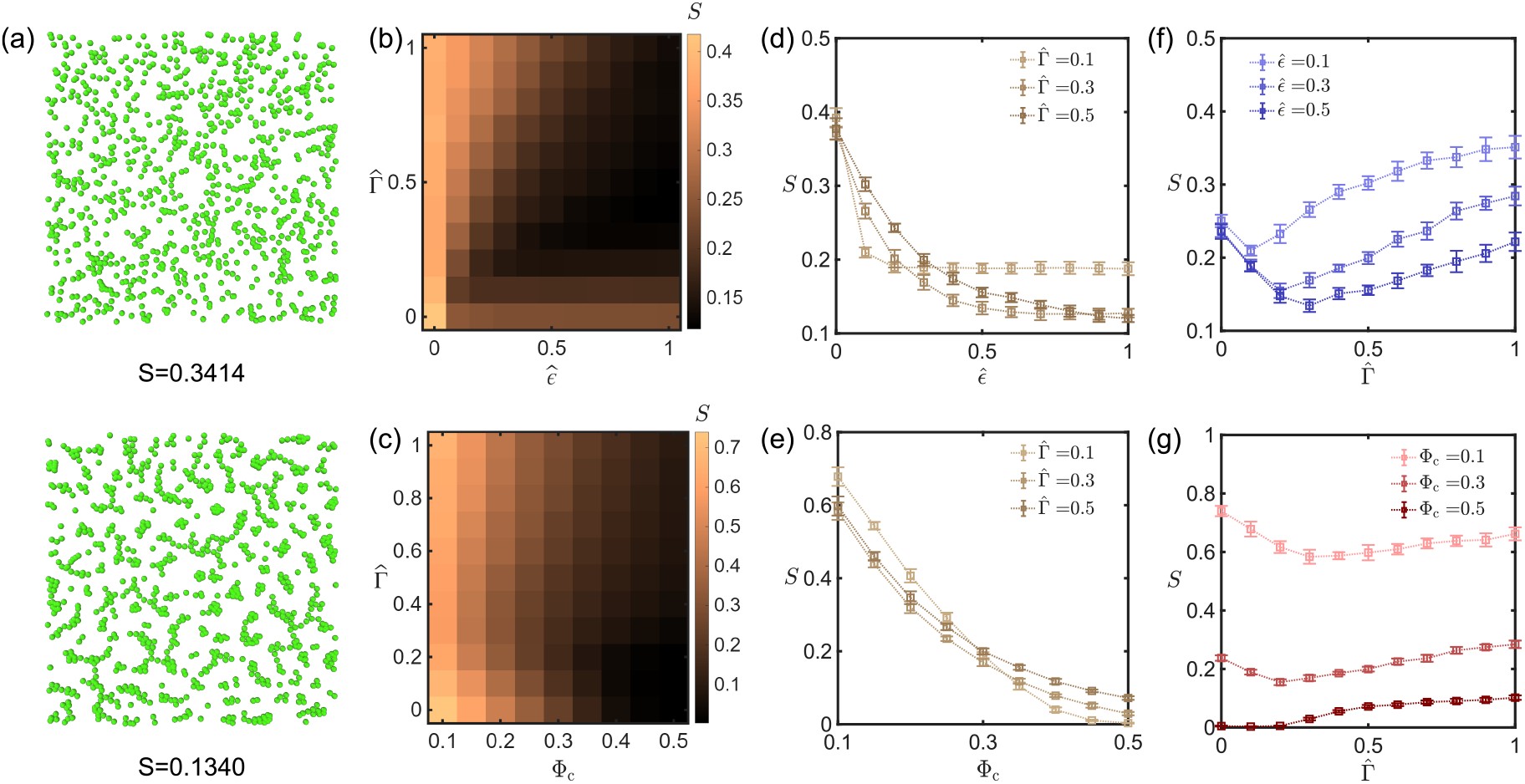
(a) Order parameter *S* quantifies the degree of self-assembly. Two snapshots in the left represents structures with *S* = 0.3414 (top), and 0.1340 (bottom) respectively. Green particles represent cells, and gels are not shown. (b–c) Phase diagrams showing the dependence of *S* on adhesion strength 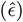, motility strength 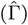, and packing fraction (*ϕ*_c_) for a 2D system at 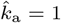 and *E*_*g*_ */E*_*c*_ = 1. (b) shows a 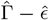 plane at *ϕ*_*c*_ = 0.3 and (c) shows a 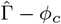plane at 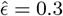. (d–g) Variation of *S* with individual parameters: (d) *S* vs. 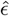at ϕ_c_ = 0.3 for different 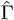; (e) *S* vs. *ϕ*_c_ at 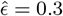 for different 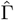; (f) *S* vs. 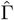at ϕ_c_ = 0.3 for different 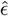; (g) *S* vs. 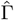at 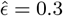 for different *ϕ*_c_.

We begin our exploration of phase-space by systematically varying the motility strength 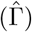, adhesion strength 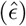, and packing fraction (*ϕ*_c_) for two-dimensional simulations keeping 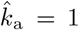 and *E*_*g*_*/E*_*c*_ = 1. Figures 2(b–c) show the dependence of *S* on these parameters in the 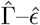 and 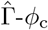 planes. As expected, *S* decreases (indicating stronger self-assembly) with increasing adhesion strength 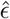 [Fig. 2(d)] and with increasing packing fraction *ϕ*_c_ [Fig. 2(e)].

In contrast, the dependence of *S* on motility strength 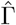 is non-monotonic. At small 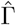, increasing motility enhances self-assembly (decreasing *S*), whereas at large 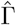, excessive motility disrupts clusters (increasing *S*) [Fig. 2(f)]. Moreover, *S* increases with 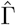 at higher densities of cells and decreases with 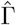 at lower densities of cells [Fig. 2(g)]. Thus, motility exerts a dual effect—it can either promote or hinder self-assembly depending on the parameter regime. Similar effect of motility in self-assembly is also observed in other active matter systems as well [29, 58].

The qualitative dependence of *S* on the three parameters are similar in three dimensions as well (Supplementary Fig. S1, Supplementary Movie 1). So, the remainder of our analysis focuses on two-dimensional systems for ease of computation.

### 3.2 Interaction of motility and adhesion in cellular self-assembly

Next, we investigate what leads to this two-way effect of motility in self-assembly and what sets the optimum motility for self-assembly. We observe the dynamics of self-assembly (Supplementary Movie 2) for the simulations with same strength of adhesion and packing densities of cells but for different strength of motility. When cells are non-motile (i.e. 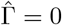), cells just locally rearrange and no specific self-assembly happens. At intermediate motility, cells can explore the simulation domain and encounter one another and adhere to each other, promoting formation of assembled structures. However, in the higher level of 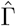, although the self-assembly happens the cells are too motile and the assembled structures become unstable.

To disentangle the two competing roles of motility in self-assembly, we quantify the number of adhesive bonds formed and broken in each simulation, defining two cells as bonded if their separation is below *r*_c_. Both quantities increase with 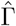, consistent with the observed two-way behavior of motility [Fig. 3(a–b)]. However, they do so in fundamentally different ways: the number of bonds formed grows steadily with increasing 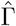, whereas the number of broken bonds remains negligible until a threshold motility Γ^∗^ is reached, after which it rises sharply. We therefore define Γ^∗^ as the minimum motility required to disrupt an adhesive cell–cell bond [Fig. 3(c), Section 5.1.3 of Methods]. We further define an optimal motility Γ_opt_ as the value of 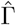 that minimizes *S* for a given 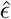 [Fig. 3(d), Section 5.1.3 of Methods], and hypothesize that Γ^∗^ and Γ_opt_ should be related. Indeed, both increase with adhesion strength 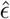, but their behaviors diverge at larger values of 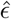 [Fig. 3(e)]. For 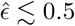, Γ^∗^ and Γ_opt_ remain approximately linearly correlated (see inset of Fig. 3(e)). As 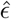 increases further, Γ_opt_ exhibits only a weak dependence on adhesion strength, whereas Γ^∗^ continues to grow. This separation indicates that, beyond a characteristic adhesion scale, increasing motility continues to break adhesive bonds but does not substantially alter the connectivity of clusters. Consistently, the dependence of the orientational order parameter *S* on 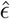 becomes markedly weaker for 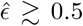 [Fig. 3(f)], indicating reduced sensitivity of cluster ordering to further increases in adhesion.

**Fig. 3.**
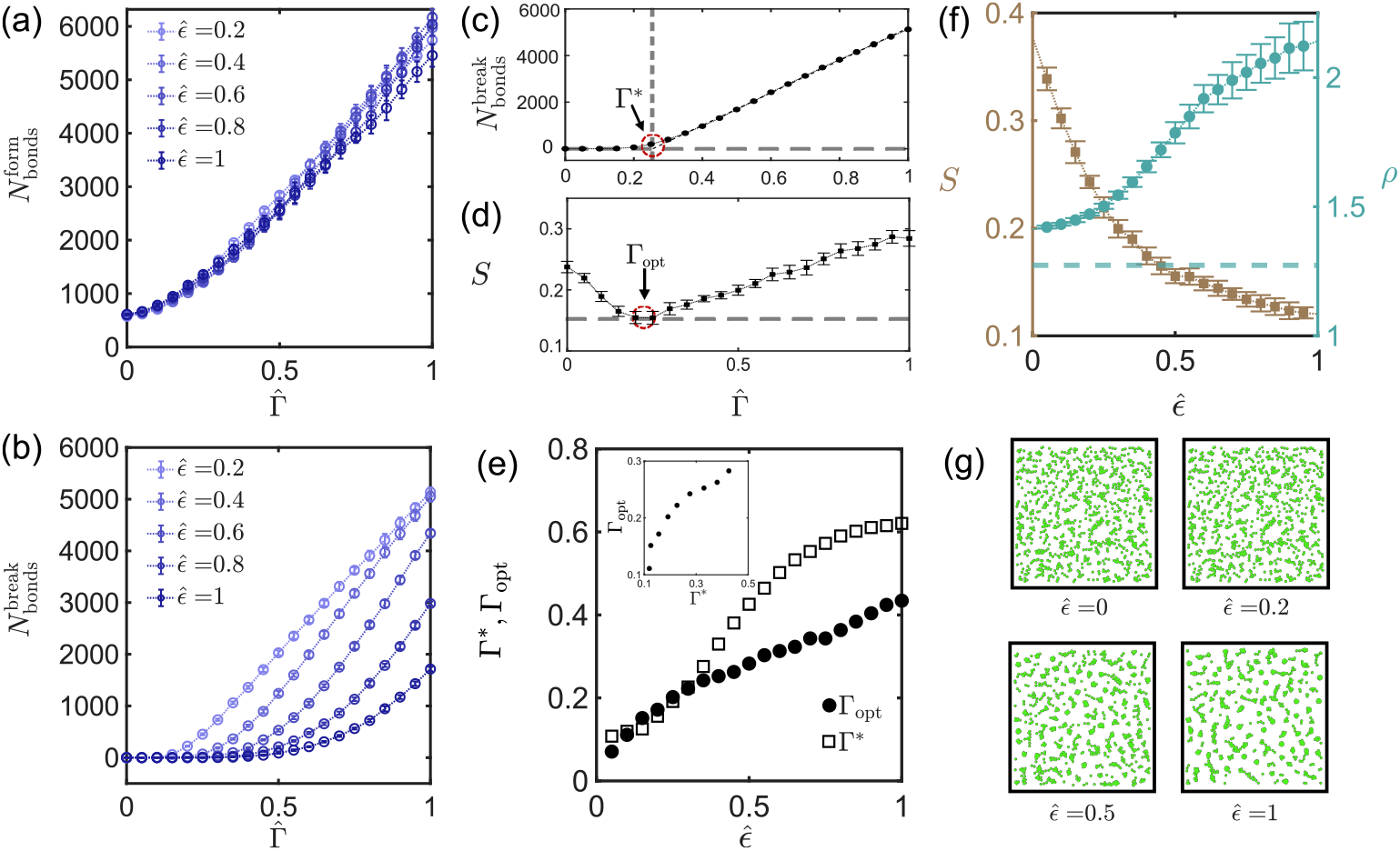
(a-b) Number of bonds (cell-cell adhesion) formed (a) and broken (b) as a function of motility strength 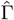for different adhesion strengths 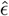. (c) Definition of Γ^∗^, the minimum motility required to break intercellular bonds. (d) Definition of Γ_opt_, the motility level at which self-assembly is maximium (*S* is minimum). (e) Variation of Γ^∗^ and Γ_opt_ on 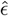. (f) Order parameter *S* (left axis) and local number density *ρ* (right axis) as functions of 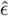. The dotted line represents number density of an isolated cell (i.e. 4/π). (g) Representative simulation snapshots showing cells (green) at 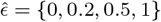 for 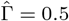 and ϕ_c_ = 0.3. Gel particles are not shown.

Why is additional motility required to break the bonds for 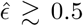 while there is no change in the number of cells in the cluster? We examine the simulation snapshots at fixed 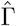 and varying 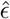. The number of clusters decreases with 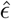 up to ∼ 0.5 and remains constant thereafter, while the clusters become increasingly compact [Fig. 3(g)]. To quantify the compactness, we calculate the average local number density of individual clusters weighted by the number of cells in each cluster [59, 60] as,

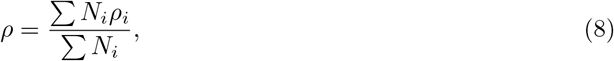

where, *N*_*i*_ is the number of cells in a cluster. *ρ*_*i*_ = *N*_*i*_*/A*_*i*_, and *ρ*_*i*_ = *N*_*i*_*/V*_*i*_ are the number density of cluster *i* for a 2D and 3D system, respectively. *A*_*i*_ and *V*_*i*_ are the non-dimensional area and volume occupied by the cluster in a 2D and 3D system respectively, with overlapping regions counted only once (Section 5.1.4 of Methods). At low values of 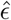, the packing fraction ρ remains close to that of an isolated sphere and exhibits only a weak dependence on 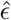. In contrast, for higher 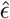, ρ increases significantly, suggesting that the clusters transition to a more compact state with substantial cell–cell overlap [Fig. 3(f), Supplementary Fig. S2(c)].

Altogether, these results suggest that our simulations show the self-assembly of cells in two regimes. For 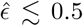, increasing adhesion primarily increases cluster size, while for 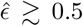, This behavior reflects the balance between adhesion and repulsion. Formation of compact clusters requires partial cell overlap, which incurs a repulsive energy cost. When adhesion is weaker than the repulsive force scale, clusters grow primarily by increasing size. Once adhesion becomes comparable to this repulsive scale, partial overlap becomes increasingly favorable, enabling denser assemblies. The value 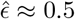 therefore represents a force-balance crossover rather than a sharp transition.

A similar analysis was carried out for a three-dimensional system as well, and the observations are consistent with that of our two-dimensional results (Supplementary Fig. S3).

### 3.3 Quantifying the effect of motility in *in vitro* cell culture experiment

Next, we asked whether the simulation-based predictions are consistent with the aggregation behavior of real cells in a viscoelastic environment. A previous study [12] reported that cells form clusters whose extent increases with both cell density and intercellular adhesion, controlled by E-cadherin overexpression. This trend closely parallels our simulation results. We designed a similar experimental assay, but instead of modulating adhesion, we tuned the strength of cellular motility. Specifically, we culture a breast cancer cell line (MCF7) in non-adherent condition in a granular and porous viscoelastic 3D cell growth media comprising jammed packings of agarose granules generated using a flash solidification strategy [49](Section 5.2.1 of Methods). Prior characterizations of such microgels have reported individual particle sizes as 100 ± 40 µm, whereas, the interparticle pore spaces range from 1-20 µm suggesting a packed microgel medium similar to our simulation. Rheological measurements confirmed that microgel medium is viscoelastic, with storage modulus exceeding loss modulus on relevant timescales, and that it exhibits yield-stress behavior (Section 5.2.2 of Methods, Supplementary Fig. S4). We performed a similar oscillatory strain response test computationally (Section 5.1.5 of Methods) for a simulation box filled with gels and we found out that similarly the storage modulus is significantly more than the loss modulus in the relevant timescales (Supplementary Fig. S5).

To modulate motility, we cultured cells at two temperatures: 22°C and 37°C with atmospheric CO_2_ concentration controlled at 5% for both conditions. Using timelapse imaging (Section 5.2.4 of Methods, Supplementary Movie 3), we tracked motion with single cell resolution in 3D. The instantaneous speed distribution shifted to higher values at 37°C. Specifically, a Kolmogorov-Smirnovtest shows that the distributions are significantly different with *p* value *<* 0.0001 and the median speed changes from 4.62 µm/hr to 14.54 µm/hr [Fig. 4(e)]. We note that the speed of an isolated cell in our model corresponds to the parameter 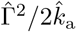, directly linking experimental results to the model parameters. Furthermore, we also calculated the mean squared displacement (MSD) and velocity-velocity corrrelation from the cellular track (Supplementary Fig. S6). We found that the velocity-velocity correalation approaches zero suggesting a loss of persistence and the MSD is in the diffusive regime with an apparent increase in diffusivity for the cells cultured in 37°C.

**Fig. 4.**
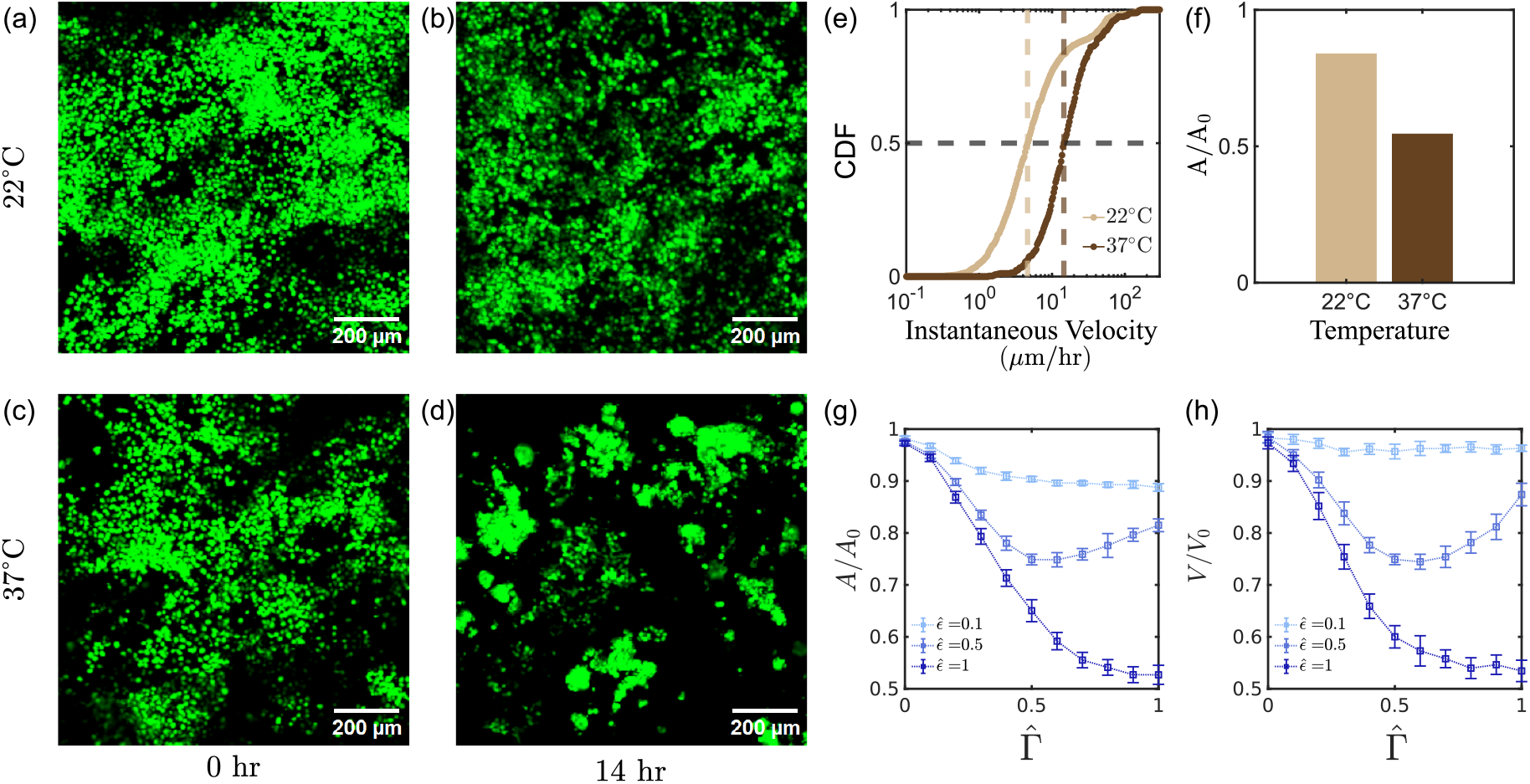
(a-d) Maximum intensity projections from micrographs of fluorescently-labelled (calcein-AM) MCF7 cells dispersed in a microgel matrix, captured at both 0 hour and post 14-hour timepoints, across both (a and b) low (22°C) and (c and d) high (37°C) temperatures. Scalebars denote 200 µm. (e) Cumulative distribution of all instantaneous speeds recorded for individual cells over four hours of timelapse imaging across both low (22°C) and high (37°C) temperatures. The horizontal dotted line represents CDF=0.5 and the verticle dotted line represents the median speed for the respective temperatures. (f) Quantification of area fractions occupied by MCF7 cells and clusters from max-projected confocal micrographs, represented as 14-hour timepoint values normalized against the 0-hour timepoint values. (g) Ratio of area occupied by the cells at the beginning and end of the 2D simulation as a function of 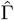 for varying 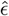 for simulations done with ϕ_c_ = 0.3, 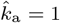 and *E*_*g*_ */E*_*c*_ = 1. (h) Ratio of volume occupied by the cells at the beginning and end of the 3D simulation as a function of 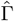 for varying 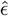 for simulations done with ϕ_c_ = 0.3, 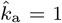 and *E*_*g*_ */E*_*c*_ = 1.

How does motility influence the self-organized clustering of these cancer cells? Over a 14-hour experiment, slower motion aberrantly affects self-organized aggregation. While MCF7 cells maintained under 22°C remain largely dispersed, cells maintained under 37°C collectively organize into aggregates [Fig. 4(a)–(d), Section 5.2.4 of Methods]. Although the imaging resolution prevents segmentation of individual cells—and thus we cannot compute the metric S, we quantified the total area occupied by cells at the beginning and end of the experiment [Fig. 4(f)]. In both conditions, the occupied area decreases over time. This reduction is significantly greater at 37°C, demonstrating that stronger motility enhances self-assembly *in vitro*.

To directly compare the experimental results to simulation predictions, we evaluated the ratio of the final to initial occupied area and volume for the 2D and 3D simulations respectively as:

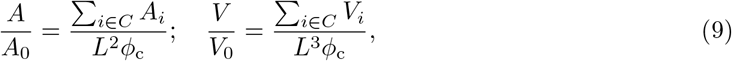

where, *A*_*i*_, and *V*_*i*_ are the areas and volumes of the individual clusters in 2D and 3D simulations respectively, *L* is the box length, and *ϕ*_c_ is cell volume fraction. We employ a Monte-Carlo integration to calculate the area and volume of the clusters. The initial area and volume are computed as *L*^2^ϕ_c_ and *L*^3^ϕ_c_ as we start from a configuration with non-overlapping cells. In simulations, *A/A*_0_ and *V/V*_0_ varies weakly with 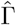 at low 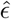, becomes non-monotonic at intermediate 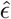, and decreases sharply at high 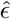 [Fig. 4(g), (h)]. We emphasize that *A/A*_0_ (or *V/V*_0_) and *S* capture different aspects of organization: *S* decreases when the number of cells in an individual cluster increases, whereas *A/A*_0_ (and *V/V*_0_) decreases primarily when clusters become compact, which occurs only at higher 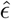 as observed in Fig. 3(f), (g). Furthermore, the indistinguishability of individual cells inside the clusters is also consistent with the high-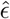 regime of our phase diagram, where we observe strong overlap and compact aggregates. Altogether, the experiments capture a regime where motility enhances self-assembly without becoming strong enough to disrupt the formed clusters, a behavior that does not appear in systems at equilibrium.

### 3.4 Effect of persistence time on self-assembly

We next examine how the persistence timescale of motility, defined as 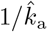, influences cellular self-assembly. To this end, we compute the order parameter *S* as a function of 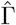 and 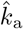while keeping 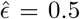 and ϕ_c_ = 0.3 fixed. We find that *S* exhibits a non-monotonic dependence on 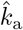, with the minimum shifting to higher 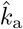 values as 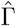increases [Fig. 5(a)–(c)]. Remarkably, all data collapse onto a single non-monotonic curve when the *x*-axis is rescaled as 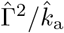 [Fig. 5(d)].

**Fig. 5.**
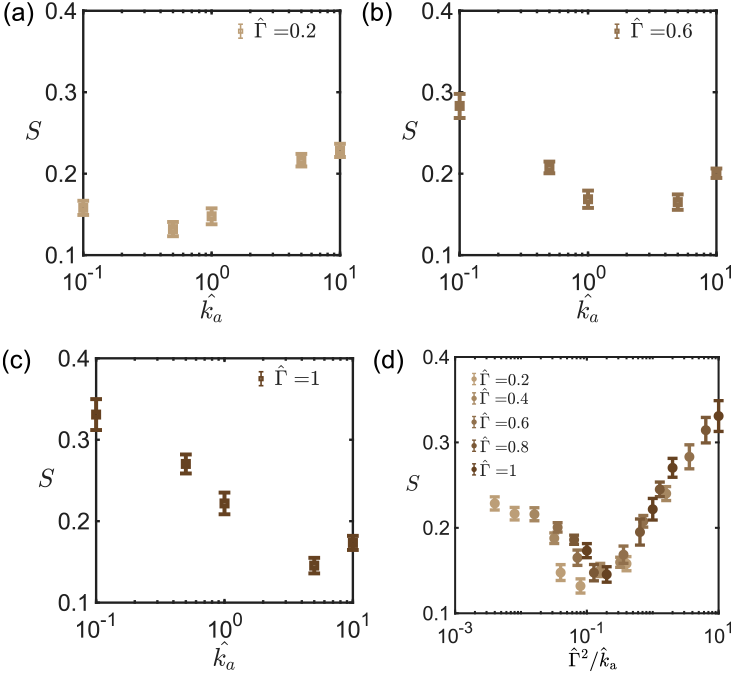
(a-c) S as a function of the inverse persistence time of motility 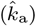 for different strengths of motility (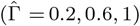, 0.6, 1), respectively for 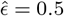 and ϕ_*c*_ = 0.3. (d) S plotted against 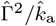 for different 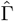 at the same 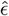 and ϕ_c_.

This rescaling has a clear physical interpretation. From Eq. 7, the zero-time value of the force–force autocorrelation function that represents the mean squared magnitude of the active force, is proportional to 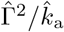. Therefore, the degree of self-assembly is primarily governed by the characteristic magnitude of the self-propulsion force. While the temporal decay of the autocorrelation does not directly control self-assembly, it influences it indirectly by modulating the effective strength of motility.

### 3.5 Effect of medium stiffness on self-assembly

In the preceding results, the background medium (gel) was assumed to have the same stiffness as the cells. We now relax this constraint to evaluate how gel stiffness influences self-assembly. We compute the order parameter *S* by varying 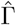and the stiffness ratio *E*_*g*_*/E*_*c*_, while keeping 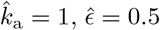, and ϕ_c_ = 0.3 fixed. We find that background stiffness significantly affects self-assembly when motility is weak. At small or zero 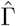, stiffer gels promote self-assembly of cells even in absence of any adhesive force 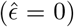 [Fig. 6(a)]. The effect persists in presence of adhesion 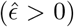 [Supplementary Fig. S7 (a)]. However, for 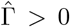, the effect of background stiffness becomes negligible irrespective of the strength of adhesion [Supplementary Fig. S7 (b),(c)].

**Fig. 6.**
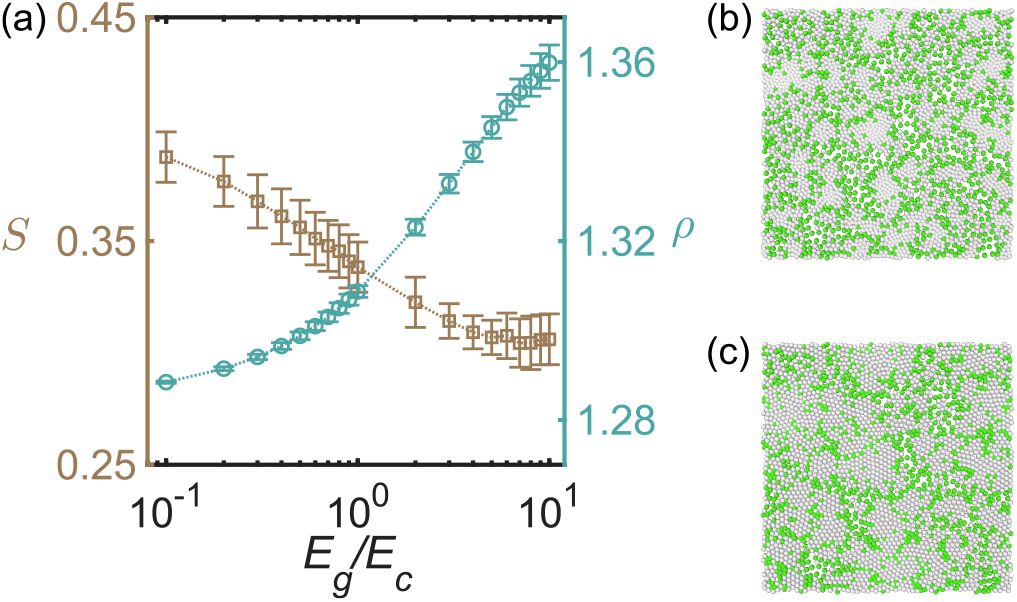
(a) S and ρ as a function of the *E*_*g*_ */E*_*c*_ for motility strength 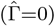 at 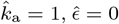 and ϕ_c_ = 0.3. (b-c) Snapshot of cellular assembly for non-motile cells (green) dispersed within gels (grey) with relative stiffness E_*g*_ */E*_*c*_ = 0.1, and *E*_*g*_ */E*_*c*_ = 10 respectively for the same simulations.

To understand this behavior, we analyze simulations with zero motility 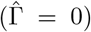. When the gel is soft (*E*_*g*_*/E*_*c*_ = 0.1), cells undergo almost no rearrangement, leading to minimal self-assembly [Fig. 6(b), Supplementary Movie 4]. In contrast, when the gel is much stiffer than the cells (*E*_*g*_*/E*_*c*_ = 10), the strong repulsive forces within the gel drive it to eliminate overlap. This compaction effectively pushes cells together even at the energy cost of their overlap promoting cluster formation even without adhesion [Fig. 6(c), Supplementary Movie 4]. This is also evident from the increasing compactness of the cluster [Fig. 6(a)]. Once motility is present, however, cells can explore space and locate neighbors autonomously, making this gel-induced “mechanical push” redundant for assembly of cells.

### 3.6 Effect of cell-cell adhesion and background stiffness in cell dynamics

Finally, we investigate the effect of different physical parameters on cell dynamics by computing the mean squared displacement (MSD) of the cells. The primary determinant of the dynamics are the strength 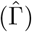 and persistence (represented by 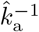) of motility. In fact, in the absence of any interaction force, the MSD of an isolated cell is only dependent on these two parameters, and can be computed analytically from Eq. 7 as,

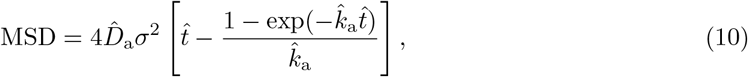

where, 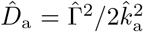 is an effective long-time diffusivity resulting from the self-propulsion, and σ is the cell diameter [Fig. 7 (a)]. This shows a ballistic behavior at short timescale 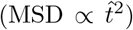 and diffusive behavior at long timescale 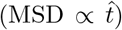, evident from the dynamics of the time exponent of MSD, 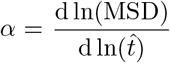 [Fig. 7 (b)]. The transition happens at the timescale of persistence of the self-propulsion force 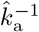.

**Fig. 7.**
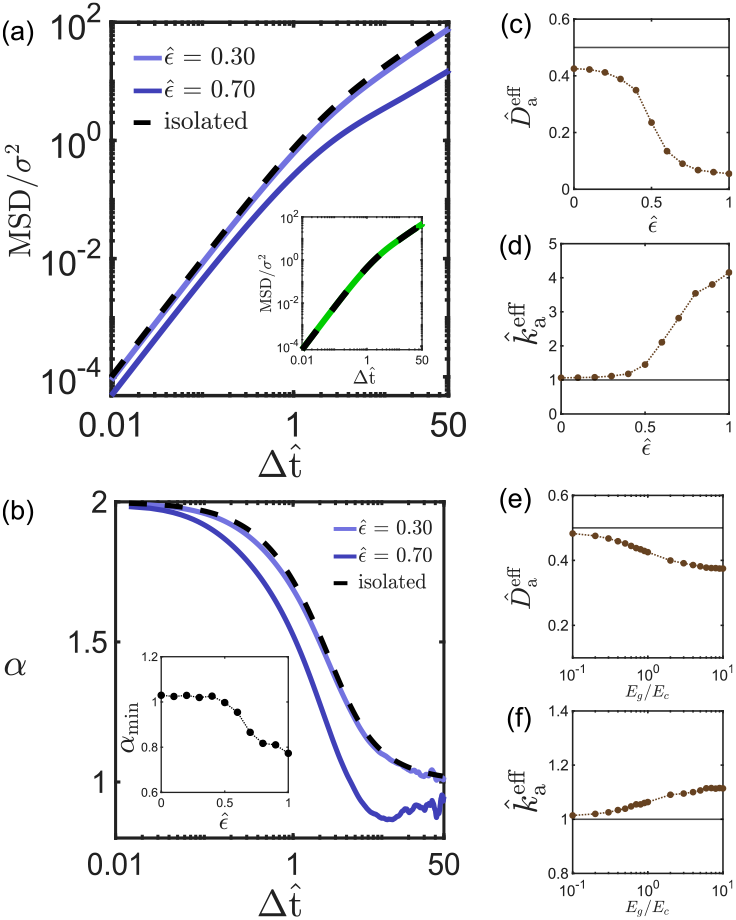
(a) Non-dimensionalized mean squared displacement (MSD/σ^2^) as a function of time for an isolated cell (derived analytically), and simulation results for 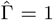 and 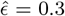 and 0.7. The inset shows the simulation MSD of the cells for 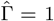 and 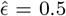, where the black dashed line represents the fit with fitting parameters 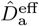and 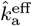. (b) The time exponent of the 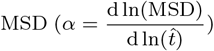, for the same three cases. The inset displays the minimum value of α from the α vs 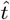 curve for varying 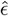, at 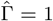. (c-d) The effective parameters 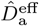 and 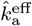 as a function of 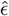 for *E*_*g*_ */E*_*c*_ = 1, 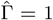, and 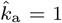, respectively. (e-f) The effective parameters 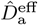and 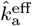 as a function of *E*_*g*_ */E*_*c*_ for 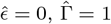, and 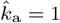. The dotted lines represent 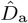 and 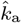for an isolated cell propelling with 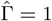, and 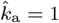.

How does this behavior change in presence of interaction with other cells and the background gels? Although the MSD reduces, we can still fit it with Eq. 10 with a modified diffusivity 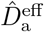and inverse persistence timescale 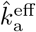 [Fig. 7(a) inset, Supplementary Fig. S8]. Specifically we asked how these quantities, which are a primary function of 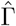 and 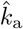 depend on strength of adhesion and background stiffness. Both quantities depend strongly on adhesion: 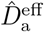 decreases and 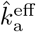 increases progressively with 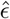 [Fig. 7(c)–(d)]. Even when adhesion is absent, 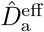 and 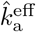 deviate from the isolated-cell prediction, suggesting an influence from the surrounding gel. To test this, we vary the background stiffness *E*_*g*_*/E*_*c*_ while setting 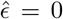. We find that stiffer gels further reduce 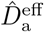 and increase 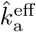 [Fig. 7(e)–(f)]. The only qualitative difference we observe from the behavior of an isolated cells is that for higher adhesion, an intermediate sub-diffusive regime emerges before long-time diffusion is recovered [Fig. 7(b)]. Notably, this behavior is observed at 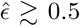, the regime of compact cluster formation [Fig. 7(b) inset]. This behavior is also observed in other active matter system with interactions such as active brownian particles with repulsive interaction [61] and polymers subject to active fluctuations [62].

Altogether, these observations suggest that presence of cell-cell adhesion as well as the interaction with the background gels reduces the net displacement as well as the persistence timescale of the cells. Physically, adhesion promotes the formation of multicellular aggregates. These assemblies are larger entities than individual cells and comprise many cells whose self-propulsion vectors are oriented in different directions, leading to partial cancellation of active forces. Consequently, both the net displacement and the effective persistence of motion are reduced relative to isolated cells. Similarly, a stiffer background gel increases mechanical resistance, making it harder for cells to migrate resulting into loss of persistence as well as overall displacement.

## 4 Conclusion

In this article, we present an agent-based computational framework, where cells are represented as soft self-propelling particles adhering to each other submerged in a passive viscoelastic environment (Fig. 1). We utilize this framework to study how the density, strength of adhesion, strength of motility, persistence time, and relative stiffness with the background determines the self-assembly of cells (Fig. 2, 5, 6). We specifically focus on the interplay of adhesion and motility in the self-assembly (Fig. 3). Further, we experimentally validate our predictions using self-aggregating cancer cells cultured in a granular microgel matrix, and discover a behavior consistent with the high-adhesion region of the phase diagram (Fig. 4). Finally, we show that the adhesive interaction with other cells and contact interaction with the background medium reduce both the strength and persistence of the motiliy of the cells (Fig. 7).

Our simulations reveal several mechanistic predictions regarding how motility and adhesion jointly regulate self-assembly. Most notably, motility plays a dual role. On one hand, active motion enables cells to explore their surroundings, encounter neighbors, and promote aggregation. On the other hand, the same motion can destabilize existing contacts and fragment clusters. Cluster formation therefore emerges from a dynamic balance between motility-driven assembly and motility-driven disruption, naturally giving rise to a non-monotonic dependence of clustering on motility strength. Although motile cells in our simulations form regions of enhanced local density, this behavior is fundamentally distinct from motility-induced phase separation (MIPS). In classical MIPS, phase separation can occur purely from self-propulsion even in the absence of attractive interactions. In contrast, we observe negligible assembly when adhesive interactions are removed, irrespective of motility strength (Supplementary Fig. S9). Thus, motility alone is insufficient; adhesion is necessary for aggregation. For this reason, we noted the observed behavior more precisely as *motility-assisted* phase separation, where motility enhances and regulates clustering but does not independently drive it. The strength of adhesion also exhibits a two-stage influence on structure formation. At lower adhesion strengths, cells form stable yet relatively loose multicellular aggregates, whereas increasing adhesion progressively compacts these clusters. This transition results from the competition between attractive and repulsive forces and is a direct consequence of our force-field design. Unlike molecular-style potentials (e.g., Lennard–Jones–type interactions), where attraction and repulsion are intrinsically coupled, our model treats them as independent physical mechanisms. Repulsion arises from mechanical stiffness and contact deformation, while adhesion reflects short-range binding. This decoupling enables independent control of these effects and produces regimes of cluster growth and compaction that would not be accessible in conventional active Brownian particle models, thereby providing a more biologically relevant model of cellular self-assembly.

We show that our simulation is qualitatively consistent with *in vitro* cell culture experiments from earlier reported results as well as new results reported here. Moreover, we would like to note that all the physical parameters are independent in our model. However, in reality they are often coupled. For example, cell migration is often modulated by the mechanical properties of the background and the strength of adhesion and motility are often moduled together by external stimulus. In future, a dedicated experimental work is needed to construct an experimental twin of the phase diagram we present in this manuscript, which will set the basis for an iterative process to reflect the coupling between the physical parameters in our model. “On the other hand, a more detailed model of the background matrix is required to account for the fibrous network structure and mechanochemical feedbacks present in extracellular matrix of *in vivo* systems.” We also note that, in this study, we only focus on the self-assembly of cells initially dispersed at random. The same framework, however, can be extended to simulate the evolution of preassembled tissues in soft background, such as those generated by 3D bioprinting [50, 63]. Specifically, the model could serve as a platform for designing tissues with switchable morphologies by tuning biophysical parameters. Thus, the framework not only provides insight into how physical interactions shape cellular self-assembly, but also lays the groundwork for the integrated design of engineered tissues that are indeed printable in soft environments.

## 5 Methods

### 5.1 Computational Methods

#### 5.1.1 Simulation setup

For all the simulations, we constructed periodic boxes with length of *L* cell diameters. For a simulation with packing fraction *ϕ*_*c*_, we calculate the number of cells 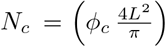, and 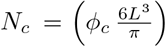 for 2D and 3D respectively. We placed the cells inside the box using Random Sequential Addition (RSA) [57] to avoid any initial overlap of cells. This algorithm is suitable for us because we vary ϕ_c_ in a range less than the maximum packing fraction for RSA. Further, we calculate the number of gels required to make an almost packed structure by setting *ϕ*_*c*_ = 1, resulting in 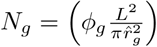 and 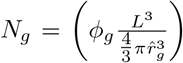, for 2D and 3D respectively, where 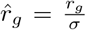. We also ensured the gels don’t overlap with cells.

After setting up initial simulation box, we evolve the coordinates of the particles using Eq. 3 using an explicit Euler scheme with timestep 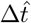.

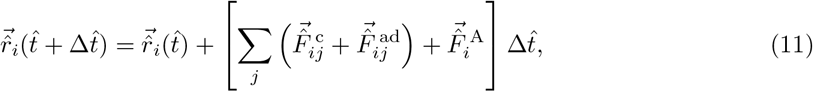

where, the contact forces 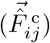 and adhesion forces 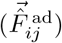, for each pair of particles were calculated using Eq. 4, and Eq. 5. The active force 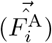 in our model follow Eq. 6 and evolves as:

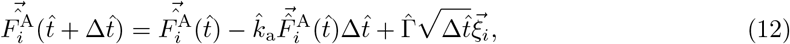

where, 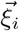is a vector with it’s all components taken from normal distribution 𝒩 (0, 1).

For all the simulations reported in the manuscript, we use *L* = 50 for 2D, and *L* = 15 for 3D. For all simulations we use a stepsize of 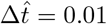. We run the simulations till *T* = 10 for all results except for calculation of MSD, where we run the simulations till *T* = 100. All the errorbars reported in the manuscript represent standard deviation from 10 independent runs.

#### 5.1.2 Identification of Clusters

To identify clusters, we first created an adjacency matrix by recognizing pairs of cells with intercellular distance less than 1 +δ cell diameters, where 2δ is the width of the attractive force-field representing adhesion (Fig. 1(b), Eq. 5). Next, we used a custom MATLAB function [64] to find the cluster of cells from the adjacency matrix.

#### 5.1.3 Calculation of Γ^∗^ and Γ_opt_

We computed the difference between adjacency matrices (A) at successive time steps, 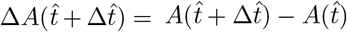. Entries with Δ*A* = +1 correspond to the newly formed bonds, whereas Δ*A* = −1 indicate broken bonds. We calculate the total number of bond formation events, 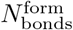 by summing all positive entries of ΔA over time as 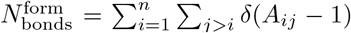, and similarly, the number of bond rupture events 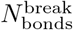 is calculated by summing over the negative entries as 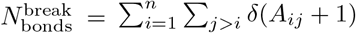.

The minimum motility required to break the bond of a particular strength adhesion 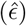 is denoted as the Γ^∗^. For each value of 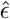, we analyze the dependence of 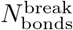 on 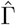. In the high motility regime, the terminal section (i.e. 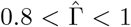) of the 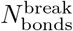 vs 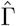 shows a consistent linear behavior across all values of 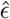. Thus, we isolated this regime and fitted using a least squares linear model of the form: 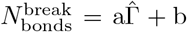, where a and b are the linear fitting parameters. The Γ^∗^ is determined by x-intercept of the fitted line yields, 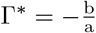[Fig. 3(c)].

We define Γ_opt_ as the motility at which self-assembly is maximum (S is minimum), as shown in Fig. 3(d). To quantify this, we fit 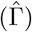 with a smooth spline using the *smoothspline* function of MATLAB. We report Γ_opt_ as the location of the minimum of the interpolated smooth spline.

#### 5.1.4 Calculation of area and volume of a cluster

To calculate the effective area (in 2D) of the cluster of cells, we determine the union of each cell’s individual area. Each cell is modeled as a disk of unit diameter. For a particular cluster consisting of *n* cells with positions 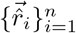, a polygonal approximation of each cell is first constructed. Specifically, the boundary of a circle of unit (cell) diameter is discretized into a set of angular points *θ* ∈ [0, 2π], and the corresponding cartesian coordinates are generated to form a closed polygon centered at 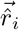. This yields a collection of *N* polygonal regions representing the spatial footprint of all cells in the cluster. The total cluster area is then obtained by computing the area of the geometric union of these individual polygonal regions using the built-in *polyshape* function in MATLAB (Supplementary Fig. S2).

An analogous procedure is adopted in 3D to estimate the cluster volume. Here, each cell is modeled as a sphere of unit diameter centered at 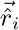, and the surface of each sphere is discretized to generate a point-cloud representation of the cell boundary. We used the built-in MATLAB function *alphaShape* to construct a polyhedron that acts as an envelope for the collection of points. The effective cluster volume is volume of this polyhedron calculated using built-in MATLAB function.

#### 5.1.5 Rheological characterization

To understand the rheological character of the gels in the simulation, we first filled our 2D simulation box of length 50 cell diameters with gel particles, and therefore considered only the contact interaction between the gels. We equilibrated the particles until *T* = 10. To perform oscillatory rheology “experiment” on the system, we impose an oscillatory shear on the system along the xy plane by applying time-dependent shear strain of the form,

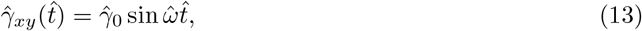

where, 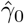 is the non-dimensionalized strain amplitude and 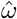 is the angular frequency of the oscillation.

This was implemented by modifying the evolution of the x-component as,

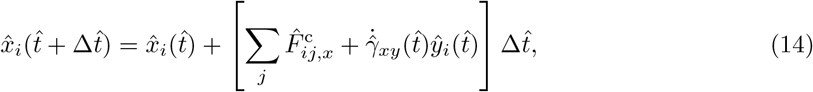

where the strain rate corresponding to (Eq. 13) is given by:

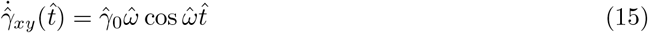

The corresponding shear stress 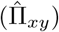, was computed using the Irving-Kirkwood formulation [65] of the stress tensor. The stress was evaluated at each simulation timestep during the oscillatory deformation as,

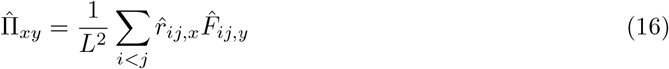

Moreover, we used Lees-Edwards boundary condition [66] to account for the strained periodic box.

For a linear viscoelastic material subjected to oscillatory shear, the stress can be expressed as a combination of two components proportional to the imposed strain and strain rate respectively (Eq. 13),

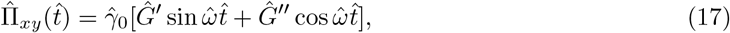

where, 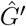 and 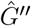 represents the non-dimensional storage and loss moduli, respectively. We obtained 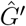 and 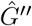 using a least square fit from the trajectories of the last five cycles of oscillation to avoid any transient effect (Supplementary Fig. S5).

### 5.2 Experimental Methods

#### 5.2.1 Preparation of 3D growth media

3D growth media is manufactured using a flash solidification method [49, 67], wherein molten agarose (40 −50°C) is rapidly injected (1 mL/sec) into a vigorously mixed bath of cold water (4°C). The rapid shearing and turbulent mixing break up the stream of agarose into micron-sized droplets, which rapidly freeze due to the temperature freezing, giving rise to solidified granules. These granules are jam-packed by centrifugation and decanting of the overlying water, yielding a gel-like mass representing the 3D matrix. This is further sterilized by brief (5 min) exposure to UV light. Following this, we wash the 3D matrix with an excess of cell culture media (DMEM supplemented with 10% FBS) by employing a process of vigorous mixing and centrifugation. Effectively, this exchanges the interstitial water with nutrient broth, thereby generating a 3D growth matrix suitable for cell culture.

#### 5.2.2 Rheological characterization

For rheological characterization, we use a shear-controlled Anton Parr 302e rheometer operated with a 50 mm roughened cone-plate measuring tool. 3D matrices prepared using the flash solidification strategy described above are subjected to oscillatory shear rheology using low amplitude (1%) strains across different frequencies to measure the shear moduli of the system. Here, the elastic storage modulus (G’) indicates a solid-like behavior, whereas, the viscous loss modulus (G”) indicates a fluid-like behavior [68]. We find that the 3D growth matrix exhibits predominantly solid-like behavior (as indicated by G’ > G”) [69]. Further, we subject the 3D matrices to unidirectional shear across varying rates while recording the shear stress response, which reveals a shear-dependent rheological behavior, characterized by fluid-like behavior under high shear regimes and solid-like behavior under low shear regimes, the transition point between which is characterized by the yield stress of the material [70].

#### 5.2.3 Cell culture and clustering assay

MCF7 cells are maintained using DMEM supplemented with 10% FBS in adherent T-25 flasks under standard culture conditions (37°C and 5% atmospheric CO_2_ concentration). Fully confluent flasks are labelled with the viability-marking green fluorescent dye calcein-AM, and subsequently harvested for setting up the clustering assay, wherein cells are dispersed within the 3D growth matrix at a final volume fraction of 0.05 (v/v). Cell-laden 3D matrices are incubated under either 22°C or 37°C under controlled 5% atmospheric CO_2_ concentration. For assessing the cell motility volumetric fluorescence timelapse imaging is performed immediately following cell seeding while maintaining the environmental conditions as described above. Next, for assessing the clustering behavior, fluorescence confocal micrographs are acquired both immediately following the assay set up, as well as approximately 14 hours post set up.

#### 5.2.4 Quantification of cell motility and area fraction

We employ custom MATLAB codes to achieve 3D reconstruction and segmentation of individual cells using the timelapse imaging data. By tracking the centroids of these cells, we quantify their instantaneous speeds between frames. We would like to note that segmentation of individual cells only works before immediately after cell seeding and before cluster formation.

For quantifying the area fraction, we acquire volumetric fluorescence confocal micrographs of the cell clustering assays at both 0-hour and approximately 14-hour timepoints. From these, we obtain maximum intensity projections for z-stacks spanning 500 microns. We perform binarization, thresholding, and subsequent quantification of the occupied area using built-in ImageJ analysis tools.

## Acknowledgements

S. Das acknowledges IIT Bombay graduate fellowship. M.S. acknowledges TIFR graduate fellowship. We acknowledge Prof. Aravind Ramanathan for generously sharing the MCF7 cells. We thank Rujula Jagadeesh for assistance with the 3D microgel preparation and characterization. S. Dutta and T.B. acknowledge funds received from Department of Biotechnology, Govt. of India vide grant no BT/PR54128/BMS/85/596/2024. S. Dutta also acknowledges seed grant from IIT Bombay vide project RD/0523-IRCCSH0-022.

## Data availability statement

The simulation software, the simulation data, and experimental images will be available on reasonable request to the corresponding authors.

## Author Contributions

S. Dutta and T.B. conceptualized the project and acquired funding. S.Das, D.P., and S. Dutta developed the simulation software. S. Das ran the simulations. S. Das and S. Dutta analyzed the simulation results. M.S. conducted the cell culture and imaging experiments and analyzed the imaging data. S. Das visualized the data. S. Dutta, S. Das, and M.S. prepared the manuscript. All the authors reviewed the manuscript.

## Supplementary information

A supplementary material file is attached with supplementary figures 1-9. Supplementary movies 1-4 are also attached separately.

