## Supplementary figure 1-9 and Movie Captions for "Biophysical Design Space for Cellular Self-assembly and Dynamics"

### 1 Supplementary Figures

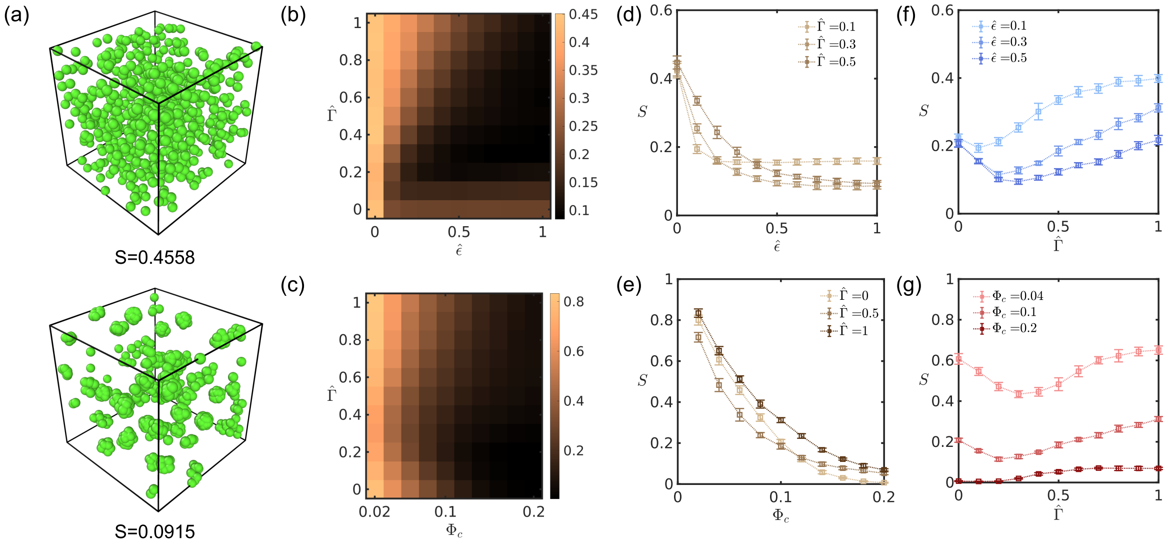


**Fig. S1** (a) Order parameter S quantifies the degree of self-assembly. Two snapshots represent structures with $S=$ 0.4558 (top), and $S = 0.0915$ (bottom) respectively. Green particles represent cells, and gels are not shown. (b-c) Phase diagrams showing the dependence of S on adhesion strength ($\hat{\epsilon}$), motility strength ($\hat{\Gamma}$) and packing fraction ($\phi_c$) for a 3D system at $\hat{k}_a = 1$ and $E_g/E_c = 1$. (b) shows a $\hat{\Gamma} - \hat{\epsilon}$ plane at $\phi_c = 0.05$ and (c) shows a $\hat{\Gamma} - \phi_c$ plane at $\hat{\epsilon} = 0.3$. (d–g) Variation of $S$ with individual parameters:(d) $S$ vs. $\hat{\epsilon}$ at $\phi_c = 0.05$ for different $\hat{\Gamma}$; (e) $S$ vs. $\phi_c$ at $\hat{\epsilon} = 0.3$ for different $\hat{\Gamma}$; (f) $S$ vs. $\hat{\Gamma}$ at $\phi_c = 0.05$ for different $\hat{\epsilon}$; (g) $S$ vs. $\hat{\Gamma}$ at $\hat{\epsilon} = 0.3$ for different $\phi_c$.

.

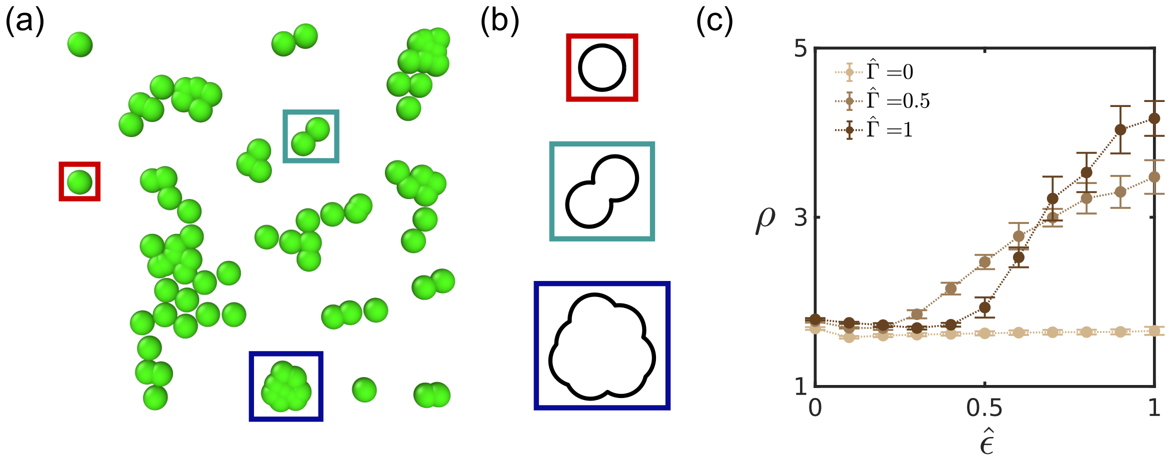


**Fig. S2** (a) A simulation snapshot with multiple clusters formed for $\hat{\Gamma} = 0.3$, $\hat{\epsilon} = 0.5$ and $\phi_c = 0.3$. (b) Enlarged views (top to bottom) showing total area occupied by a single cell, two cell and a multicellular cluster, respectively. This area is used in the calculation of $\rho$. (c) Average local packing fraction $\rho$ vs. $\hat{\epsilon}$ at $\phi_c = 0.3$ for different $\hat{\Gamma}$.

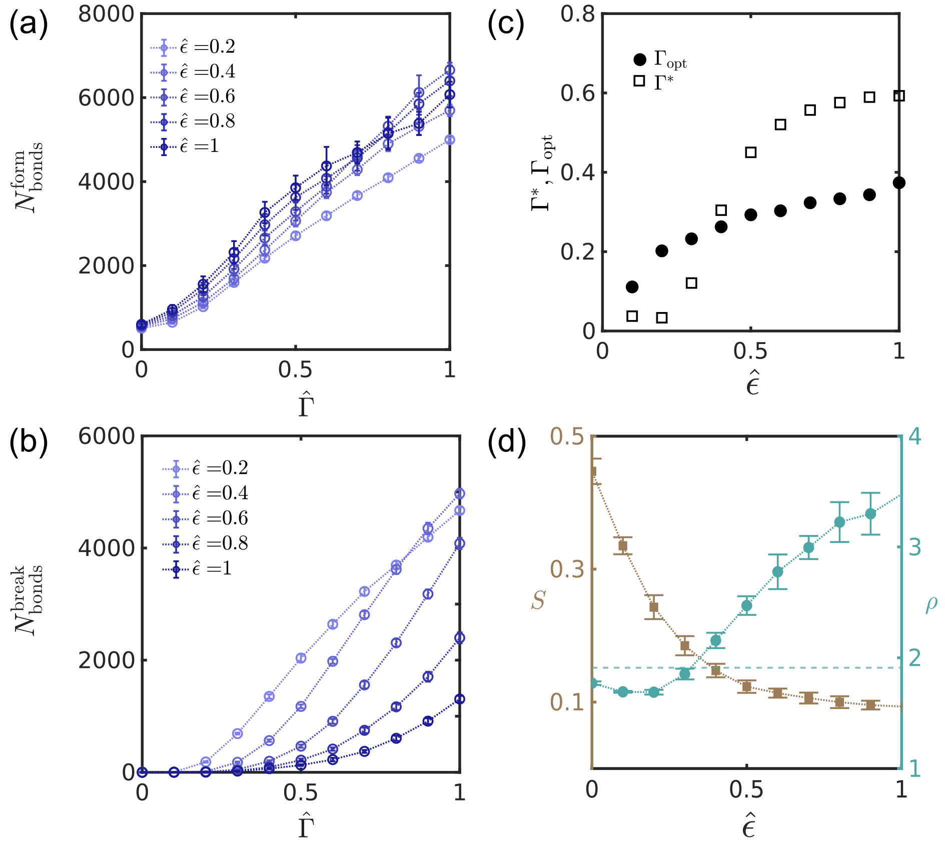


**Fig. S3** (a-b) Number of bonds (cell-cell adhesion) formed (a) and broken (b) as a function of motility strength $\hat{\Gamma}$ for different adhesion strengths $\hat{\epsilon}$. (c) Variation of $\Gamma^*$ and $\Gamma_{\mathrm{opt}}$ on $\hat{\epsilon}$. (d) Order parameter S (left axis) and local number density $\rho$ (right axis) as a function of $\hat{\epsilon}$. The dotted line represents number density of an isolated cell (i.e., $6/\pi$).

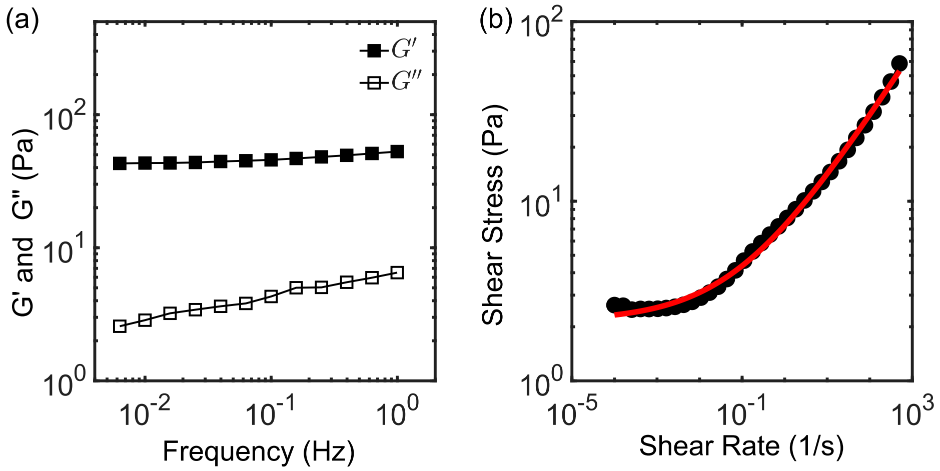


**Fig. S4** (a) Storage modulus ($G'$) and loss modulus ($G''$) of the 3D growth matrix plotted against the frequency. (b) Shear stress vs the shear rate showing experimental data (black circle) and the corresponding fit (red line) for a yield-stress behavior.

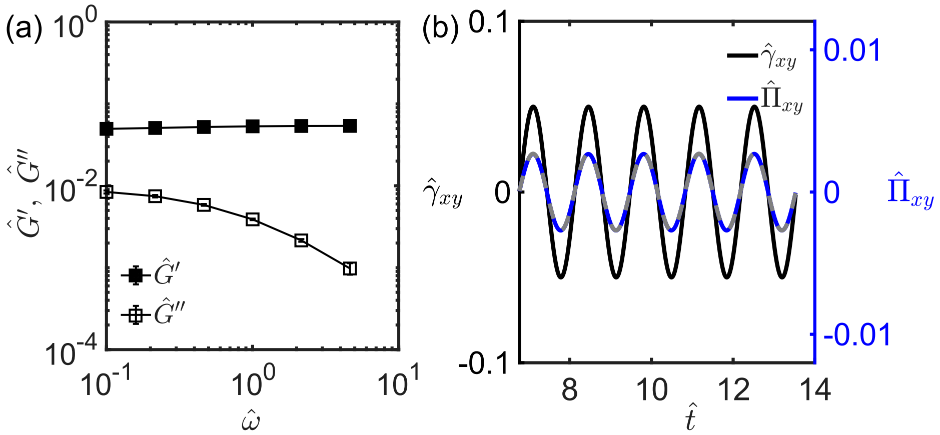


**Fig. S5** (a) Non-dimensionalized storage modulus ($\hat{G}'$) and loss modulus ($\hat{G}''$) of the gel network in the simulation box as a function of the oscillation frequency $\hat{\omega}$. (b) Non-dimensionalized strain ($\hat{\gamma}_{xy}$) and stress ($\hat{\Pi}_{xy}$) as a function of time, showing the imposed oscillatory deformation and the corresponding stress response of the gel network in the steady-state regime. The gray dotted line represents the fitted non-dimensionalized stress signal.

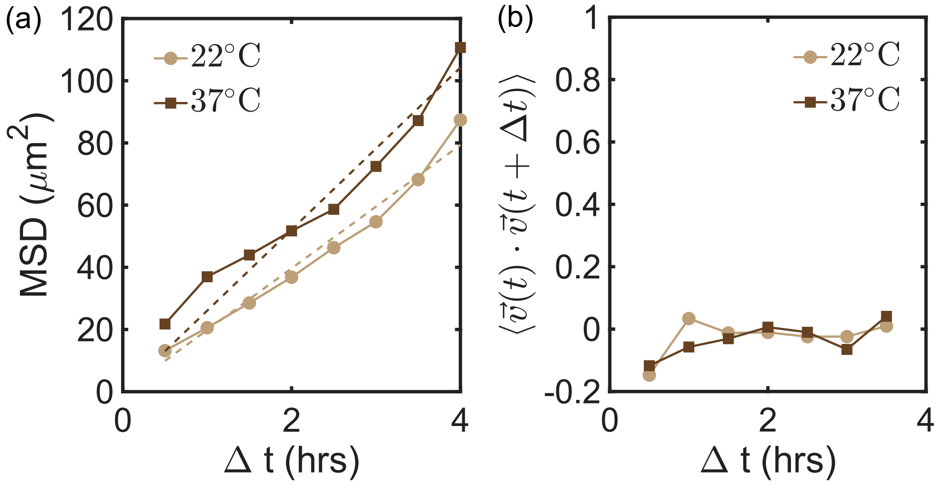


**Fig. S6** (a) MSD and (b) velocity–velocity time correlations as a function of time for cells cultured at $22^0$C and $37^0$C. The dotted lines represent the linear fits.

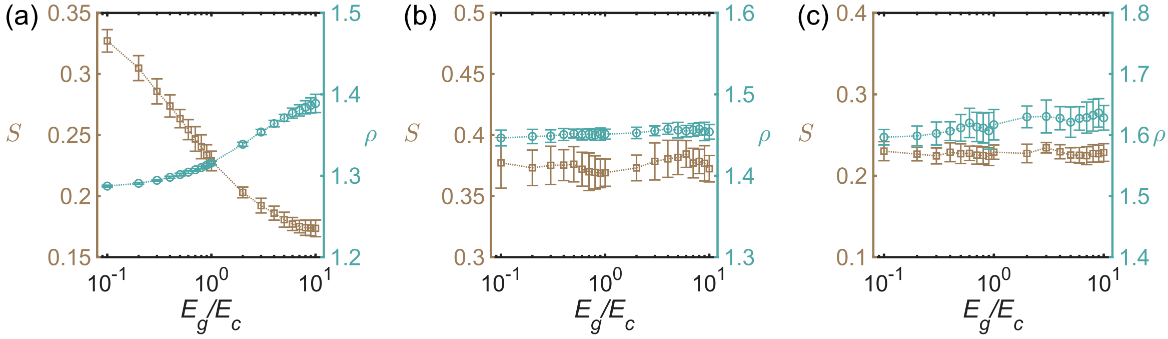


**Fig. S7** (a) Variation of S and $\rho$ with $E_g/E_c$ for $\phi_c = 0.3$ and $\hat{\epsilon} = 0.5$ at $\hat{\Gamma} = 0$. (b-c) Variation of S and $\rho$ with $E_g/E_c$ for $\phi_c = 0.3$ and $\hat{\Gamma} = 1$ at $\hat{\epsilon} = 0$ and 0.5, respectively.

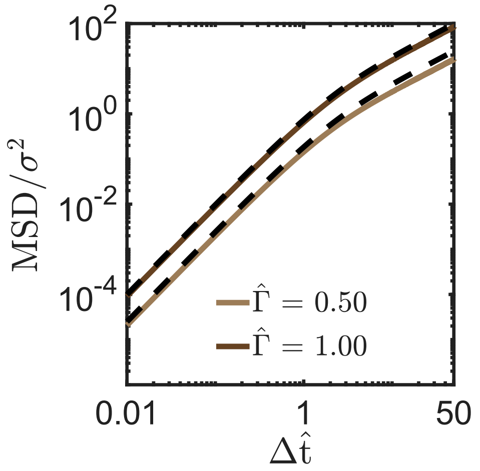


**Fig. S8** Mean squared displacement (MSD) vs time plots $\phi_c = 0.3$, $\hat{\epsilon} = 0$ and $E_g/E_c = 1$ for different values of $\hat{\Gamma}$. The dashed lines represent the fitted MSD curves obtained using fitting parameters $\hat{D}_a^{\rm eff}$ and $\hat{k}_a^{\rm eff}$.

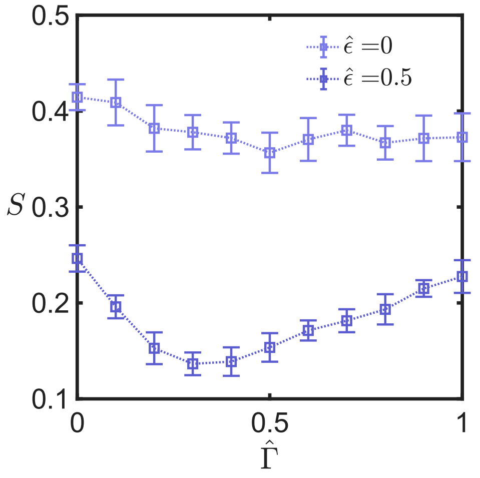


**Fig. S9** Order metric $S$ as a function of $\hat{\Gamma}$ and $\phi_c = 0.3$ for $\hat{\epsilon} = 0$ and $\hat{\epsilon} = 0.5$.

### 2 Supplementary Movie Captions

1. Movie 1: Timelapse video made from snapshots of a three-dimensional simulation with $\hat{\Gamma} = 0.5$, $E_g/E_c = 1$, $\hat{k}_{\mathrm{a}} = 1$, $\hat{\epsilon} = 0.5$ and $\phi_{\mathrm{c}} = 0.05$. Green particles represent the cells and gray transparent particles represent background (gel).
2. Movie 2: Timelapse videos made from snapshots of two-dimensional simulations for $\hat{\epsilon} = 0.5$ and $\phi_{\mathrm{c}} = 0.3$ at different motility strengths: $\hat{\Gamma} = 0,\ 0.3$ and 1. Background (gel) particles are not shown.
3. Movie 3: Timelapse videos of maximum intensity projections of MCF7 cells (fluorescently labeled with calcein-AM) dispersed within a microgel matrix at temperatures 22°C and 37°C over 4 hours with images taken in 30 min intervals.
4. Movie 4: Timelapse videos corresponding to two-dimensional simulations for $\hat{\epsilon} = 0$, $\hat{\Gamma} = 0$ and $\phi_{\mathrm{c}} = 0.3$ shown for different stiffness ratios of gels (shown in gray) and cells (shown in green): $\hat{E}_g/E_c = 0.1$(left), and 10(right), respectively.
